# Transdifferentiation is uncoupled from progenitor pool expansion during hair cell regeneration in the zebrafish inner ear

**DOI:** 10.1101/2024.04.09.588777

**Authors:** Marielle O. Beaulieu, Eric D. Thomas, David W. Raible

**Affiliations:** Molecular and Cellular Biology Graduate Program, University of Washington, Seattle, WA; Virginia Merrill Bloedel Hearing Research Center, Department of Otolaryngology - Head and Neck Surgery, University of Washington, Seattle, WA; Neuroscience Graduate Program, University of Washington, Seattle, WA; Department of Biological Structure, University of Washington, Seattle, WA

**Keywords:** zebrafish, inner ear, hair cell, regeneration, transdifferentiation Word count: 6687

## Abstract

Death of mechanosensory hair cells in the inner ear is a common cause of auditory and vestibular impairment in mammals, which have a limited ability to regrow these cells after damage. In contrast, non-mammalian vertebrates including zebrafish can robustly regenerate hair cells following severe organ damage. The zebrafish inner ear provides an understudied model system for understanding hair cell regeneration in organs that are highly conserved with their mammalian counterparts. Here we quantitatively examine hair cell addition during growth and regeneration of the larval zebrafish inner ear. We used a genetically encoded ablation method to induce hair cell death and observed gradual regeneration with correct spatial patterning over two weeks following ablation. Supporting cells, which surround and are a source of new hair cells, divide in response to hair cell ablation, expanding the possible progenitor pool. In parallel, nascent hair cells arise from direct transdifferentiation of progenitor pool cells uncoupled from progenitor division. These findings reveal a previously unrecognized mechanism of hair cell regeneration with implications for how hair cells may be encouraged to regenerate in the mammalian ear.

**SUMMARY STATEMENT:** Hair cell regeneration in the zebrafish inner ear occurs through transdifferentiation after a transient wave of supporting cell proliferation that expands the precursor pool.

## INTRODUCTION

The sensory organs of the inner ear that detect sound and head position are highly conserved across the vertebrate kingdom. The potential to regenerate these organs, however, is not as widespread. Hair cells, the mechanosensory cells of the inner ear, are particularly fragile and are vulnerable to death caused by exposure to ototoxic drugs, injury, and age-related degeneration. While mammals can regenerate hair cells at perinatal stages (Burns et al., 2012a; Mellado Lagarde et al., 2014; White et al., 2006), this ability declines rapidly after birth (Burns et al., 2012b; Cox et al., 2014; Maass et al., 2015). By adulthood, regeneration is limited in mammalian vestibular organs (Bucks et al., 2017; Forge et al., 1993; Golub et al., 2012; Kawamoto et al., 2009) and completely lost in the auditory system (Oesterle et al., 2008). As a result, hair cell death can lead to permanent auditory and vestibular deficits in humans. In contrast, many other vertebrates, including fish, amphibians, and birds, can regenerate functional hair cells throughout life (Avallone et al., 2008; Baird et al., 1996; Corwin and Cotanche, 1988; Cruz et al., 1987; Cruz et al., 2015; Harris et al., 2003; Jimenez et al., 2021; Jones and Corwin, 1996; Lombarte et al., 1993; Ryals and Rubel, 1988; Smith et al., 2006; Taylor and Forge, 2005; Weisleder and Rubel, 1992).

Zebrafish are well-known for their regenerative potential and are commonly used to study hair cell development, death, and regeneration (reviewed in Pickett and Raible, 2019; Sheets et al., 2021). In addition to inner ear hair cells, fish and amphibians have analogous hair cells in an external sensory system called the lateral line, which is used to detect changes in water flow for behaviors such as schooling and predator evasion. Much of our current understanding of zebrafish hair cell function and regeneration comes from studies of the lateral line, while zebrafish inner ear hair cells have been relatively understudied. The zebrafish inner ear remains a promising model system for studying hair cell regeneration due to its high level of conservation with the inner ear of mammals and the extensive genetic and imaging tools available for zebrafish.

Zebrafish share several conserved inner ear organs with other vertebrates: three cristae, which sense angular rotation of the head within the semicircular canals, and two otolith organs, or maculae: the utricle and saccule (Fig. 1). In mammals, the utricle and saccule sense gravity and linear acceleration while an additional structure, the cochlea, is highly specialized for hearing. Zebrafish do not have a cochlea; instead, auditory function is distributed across the macular organs, with the saccule likely playing an outsized role (Breitzler et al., 2020; Schuck and Smith, 2009). Only the utricle is indispensable for vestibular function (Riley and Moorman, 2000), but both macular organs have some capacity to respond to both auditory and vestibular stimuli (Favre-Bulle et al., 2020; Popper and Fay, 1993; Yao et al., 2016).

**Figure 1.**
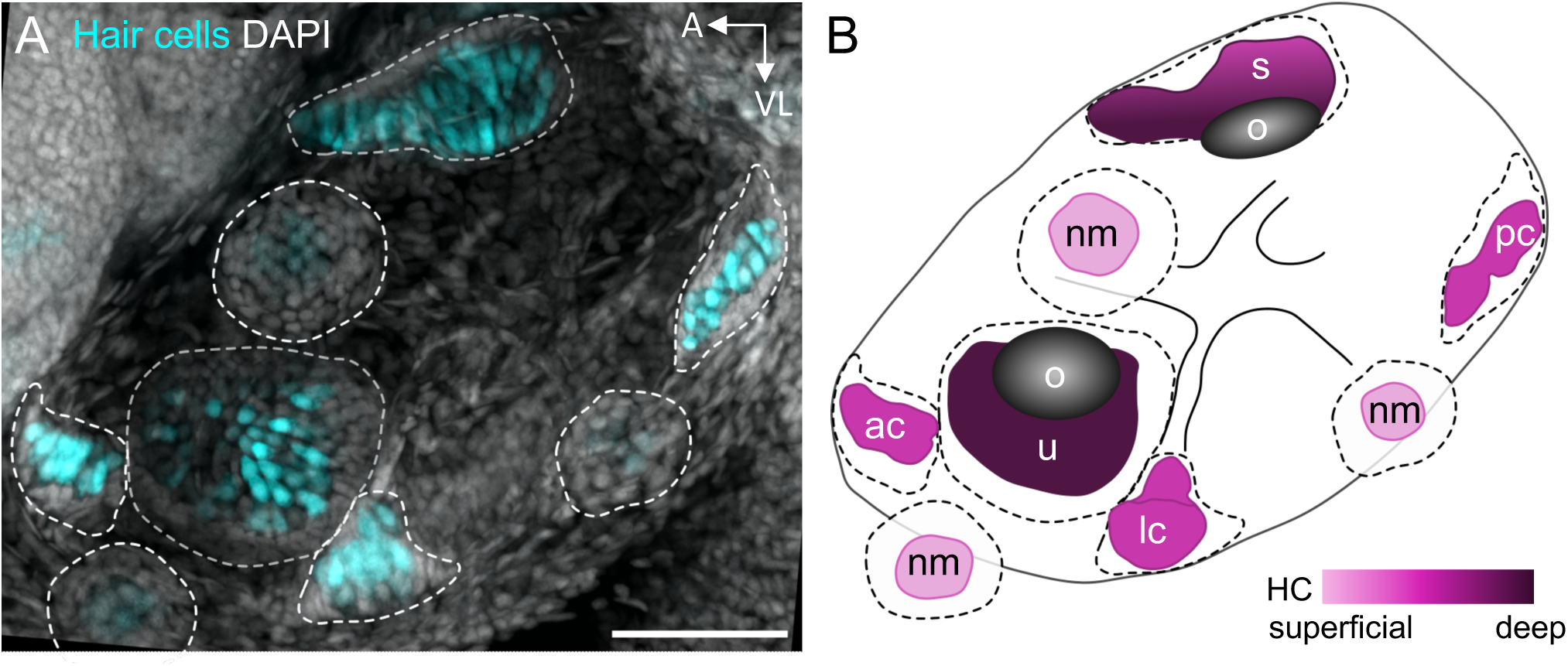
Inner ear organs of the larval zebrafish. A) Maximum intensity projection image of *Tg(myo6b:GFP)* 5dpf larval zebrafish ear. GFP-labeled hair cells are shown in cyan and DAPI-labeled nuclei are shown in grey. Dotted outlines delineate neuromast and inner ear organ boundaries. Scale bar = 50 µm. B) Diagram of 5dpf larval zebrafish ear. Color gradient indicates depth of organs where lighter colors indicate more superficial structures and darker colors indicate deeper structures. Dotted outlines delineate neuromast and inner ear organ boundaries, while color-filled areas indicate location of hair cells. ac = anterior crista, lc = lateral crista, nm = neuromast, o = otolith, pc = posterior crista, s = saccule, u = utricle.

Within a single sensory organ, hair cells can be divided into subtypes based on differences in morphology, physiology, innervation, and gene expression. Mammalian and avian vestibular organs have type I hair cells, which are flask-shaped and innervated by afferent neurons that form calyces around hair cell bodies, and type II hair cells, which have shorter bodies with long foot-like basal projections that are innervated by bouton synapse-forming afferents (Burns and Stone, 2017; Eatock and Songer, 2011). The maculae can be divided into a central striolar region and more peripheral extrastriolar regions. Zebrafish have striolar and extrastriolar hair cells analogous to those of other vertebrates (Chang et al., 1992; Jiang et al., 2017; Liu et al., 2022; Platt, 1993; Shi et al., 2023; Tanimoto et al., 2022), but they do not have the equivalent morphological and innervation dimorphism of type I/II hair cells. The cristae are also organized into central and peripheral zones with molecularly distinct hair cells (Bang et al., 2001; Haddon and Lewis, 1996; Shi et al., 2023; Zhu et al., 2021).

Hair cells are surrounded by and interspersed with supporting cells that perform many critical roles during the life and death of hair cells (Wan et al., 2013), including acting as hair cell precursors (Corwin and Cotanche, 1988; Lin et al., 2011; Lopez-Schier and Hudspeth, 2006; Millimaki et al., 2010; Weisleder et al., 1995). The mechanism by which hair cells are regenerated differs by model system, with a critical point of difference being whether precursors divide before giving rise to new hair cells. In the lateral line, nascent hair cells are added in pairs as a result of symmetric division and differentiation of supporting cells (Lopez-Schier and Hudspeth, 2006; Mackenzie and Raible, 2012; Romero-Carvajal et al., 2015; Wibowo et al., 2011). When regeneration is observed in mature mammalian vestibular organs, hair cells are added by direct transdifferentiation of supporting cells (Golub et al., 2012). A dual mechanism has been observed in the auditory organ of birds, whereby hair cells are regenerated in an initial wave of transdifferentiation followed by a later wave of asymmetric proliferation (Roberson et al., 1996; Roberson et al., 2004). Previous studies have demonstrated hair cell regeneration in the zebrafish inner ear (Jimenez et al., 2021; Millimaki et al., 2010; Schuck and Smith, 2009), with evidence for both proliferative replacement and transdifferentiation, but definitive experiments are lacking. The transdifferentiation hypothesis is supported by recent single cell and nucleus RNA-seq data, which suggest that the inner ear does not have a clear mitotically cycling supporting cell population as is seen in the lateral line (Baek et al., 2022; Lush et al., 2019) and instead show a substantial transition state population during regeneration which shares gene expression aspects of both hair cells and supporting cells (Jimenez et al., 2022).

Here, we describe a mechanism of hair cell regeneration in the zebrafish inner ear in which supporting cell proliferation in response to hair cell death is not directly coupled with the differentiation of regenerating hair cells. First, we used transgenic zebrafish lines to determine the timecourse of hair cell addition during larval zebrafish development. We found that hair cells are added at a linear rate corresponding to fish growth and that few hair cells are removed due to hair cell turnover during this time. Both hair cell subtypes of the cristae are added at equivalent rates, with some cells converting from peripheral to central subtype over time to maintain organ patterning. When crista hair cells are ablated, hair cell numbers recovered relatively slowly over the course of two weeks, and central-type hair cells are produced at an increased rate to recover proper organ patterning. We provide evidence that most regenerating hair cells are formed by transdifferentiation. We find that ablation causes an initial burst of supporting cell division, but new hair cells are not differentially derived from this dividing population. Rather, hair cell numbers recover during regeneration due to a transient increase in progenitor pool size.

## RESULTS

### Zebrafish inner ear sensory patches grow constantly during the larval stage

Sensory patches in the fish inner ear add new hair cells continuously throughout the animal’s life (Bang et al., 2001; Corwin, 1981; Corwin, 1983; Higgs et al., 2002; Higgs et al., 2003). To distinguish hair cell regeneration from addition during growth, we first quantified the rate of hair cell addition under homeostatic conditions. We examined the larval stage, during which the inner ear organs become functional and remain superficial enough for imaging in intact fish. Variation in environmental factors greatly affect fish growth: standard length (SL), a measurement from the snout tip to the caudal peduncle, becomes a better indicator of developmental stage than time after 5 days post-fertilization (dpf) (Parichy et al., 2009). The larval stage begins at 72 hours post-fertilization and continues until 30-45 dpf when the fish are 11 mm SL. The utricle is formed and functional by 4 dpf (Mo et al., 2010; Riley and Moorman, 2000), and contains both striolar and extrastriolar type hair cells. The cristae do not become functional until later on when the larvae are 8 mm SL, around 30 dpf (Beck et al., 2004), when the semicircular canals are large enough to allow for adequate fluid flow to stimulate hair cells. The cristae, however, are formed by 5 dpf and contain both central and peripheral hair cell subtypes (Bang et al., 2001; Haddon and Lewis, 1996; Shi et al., 2023; Zhu et al., 2021).

To determine the baseline rate of hair cell addition in the zebrafish inner ear, we used a *Tg(myo6b:nls-Eos)* (Cruz et al., 2015) transgenic zebrafish, which expresses the photoconvertible protein Eos in hair cell nuclei. In both cristae and utricle, hair cells were added at a steady rate (Fig. 2A-C). In anterior and lateral cristae, hair cells were added at a rate of 19.14 ± 1.33 per mm SL and 20.30 ± 1.26 per mm SL, respectively, with no significant difference between these rates (Fig. 2D). In the utricle, hair cells were added at a rate of 49.96 ± 4.3 per mm SL (Fig. 2E). Among the cristae, the lateral crista is the earliest to form and is slightly larger than the anterior and posterior cristae at the beginning of the larval stage. This size discrepancy is maintained over time, while the anterior and posterior cristae remain similar in size (Figs 2D, S1). Due to its similarity in size to the anterior crista and depth in larger fish, the posterior crista was not a focus of subsequent experiments. These results indicate that hair cells are added at a consistent rate in each of the sensory organs as larvae grow.

**Figure 2.**
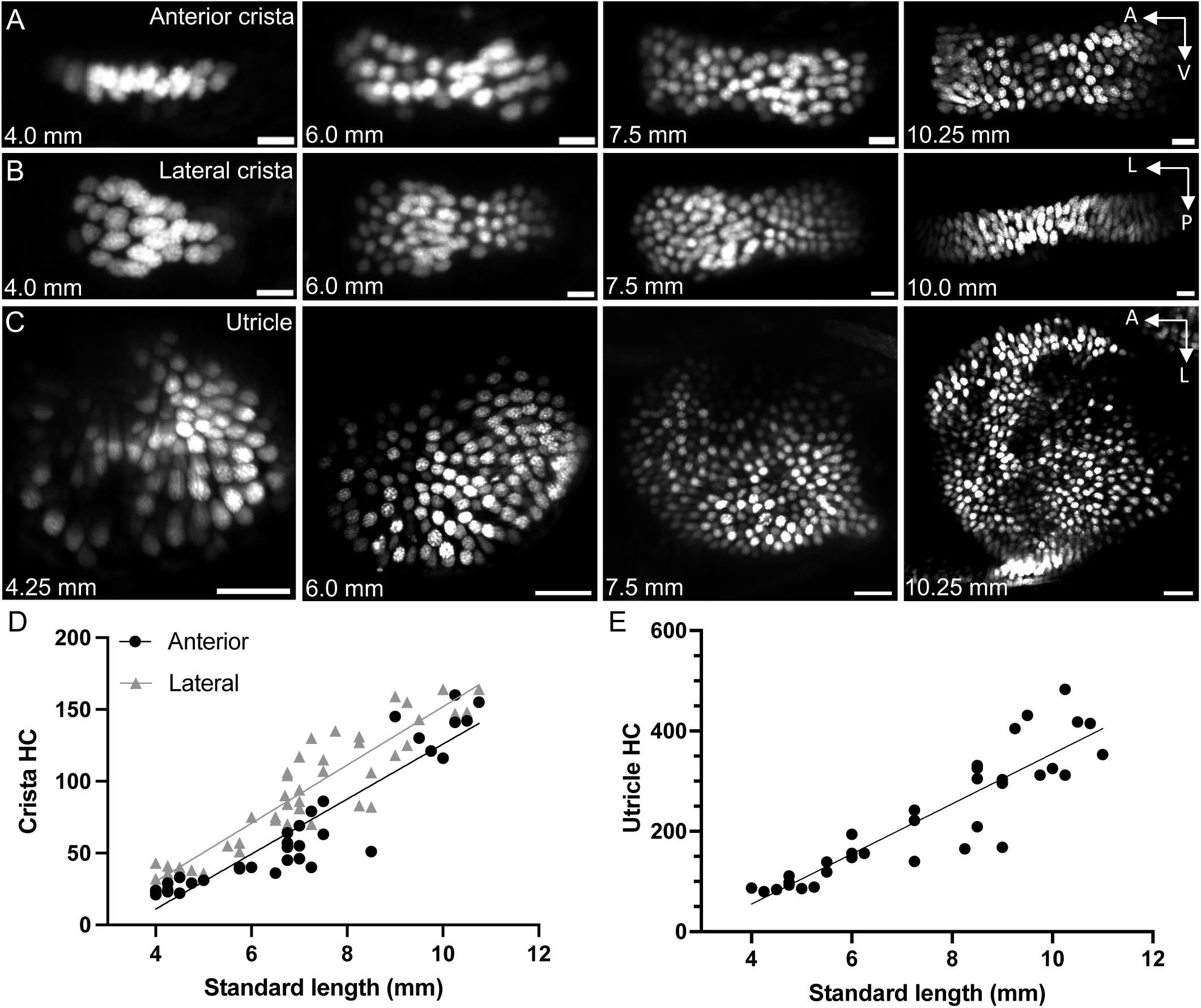
Addition of hair cells during larval zebrafish growth. A) Maximum intensity projections of *Tg(myo6b:NLS-Eos)* anterior crista hair cells at standard lengths 4.0 mm, 6.0 mm, 7.5 mm, and 10.25 mm. Scale bar = 10μm. B) Maximum intensity projections of lateral crista hair cells at standard lengths 4.0 mm, 6.0 mm, 7.5 mm, and 10.0 mm. Scale bar = 10μm. C) Maximum intensity projections of utricle hair cells at standard lengths 4.25 mm, 6.0 mm, 7.5 mm, and 10.25 mm. Scale bar = 20μm. D) Quantification of hair cell number in the anterior and lateral cristae across the larval stage of development. Anterior crista data points are represented by black circles (n = 35), while the lateral crista is represented by grey triangles (n = 47). Each data point represents one ear from one fish. Anterior crista slope = 19.14 ± 1.33 HC per mm SL, R^2^ = 0.862. Lateral crista slope = 20.30 ± 1.26 HC per mm SL, R^2^ = 0.853. Simple linear regression indicates no significant difference between these two slopes (p = 0.529). E) Quantification of utricle hair cell number across the larval stage (n = 34). Linear regression of utricle hair cell number slope = 49.96 ± 4.28 HC per mm SL, R^2^ = 0.812.

### Little hair cell turnover occurs in the developing inner ear organs

Hair cells regularly turn over in the adult zebrafish lateral line, with a half-life of approximately one week (Cruz et al., 2015). Studies from birds and mice suggest that the rate of turnover varies across species (Bucks et al., 2017; Goodyear et al., 1999; Jørgensen and Mathiesen, 1988; Kil et al., 1997). To determine the rate of turnover in the zebrafish inner ear, we again used the *Tg(myo6b:nls-Eos)* line.

Eos exhibits an irreversible green to red photoconversion upon exposure to UV light. Larval fish were placed under UV light for 10 minutes at 8 dpf (SL 4.0 – 4.5) and fixed and imaged either immediately after photoconversion (Fig. 3A, D, G) or following one week of growth (Fig. 3B, E, H). Hair cells that are added post-photoconversion can be identified by the absence of photoconverted Eos in their nuclei, while older cells retain the converted Eos signal. The anterior crista, lateral crista, and utricle showed no significant decrease in photoconverted hair cell nuclei over the course of one week (Fig. 3C, F, I).

**Figure 3.**
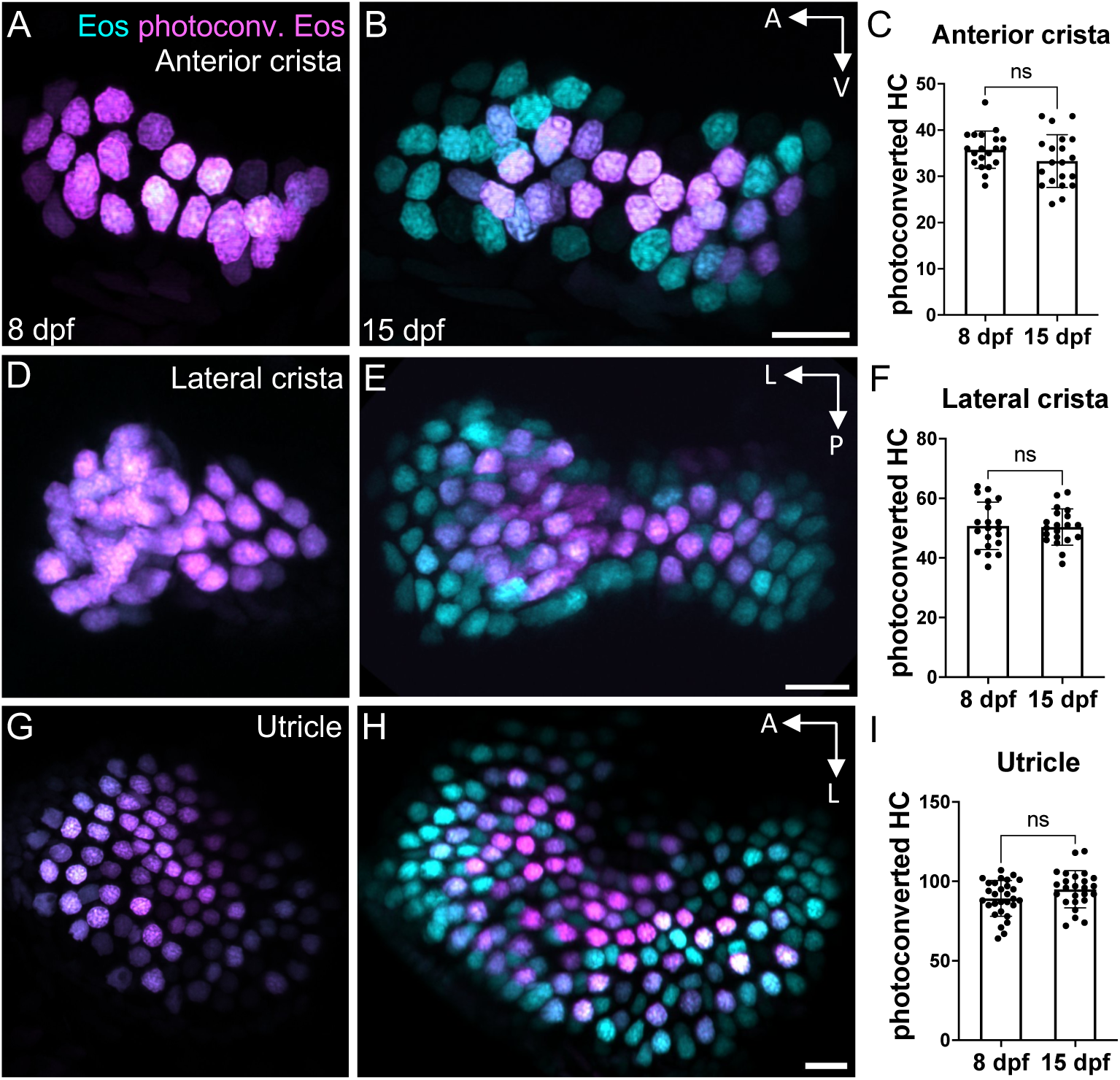
Little hair cell turnover occurs in the larval zebrafish ear. A-B) Representative maximum intensity projection images of *Tg(myo6b:NLS-Eos)* anterior cristae A) immediately post-photoconversion at 8 days post-fertilization (dpf) or B) one week post-photoconversion at 15 dpf. HC that were photoconverted retain photoconverted (magenta) Eos signal while new HC have unconverted (cyan) Eos only. C) Quantification of anterior crista photoconverted hair cells at 8 and 15 dpf (n = 20 control, 20 ablated). D-F) Analogous results for the lateral crista (n = 20, 20) and G-I) for the utricle (n = 29, 25). Unpaired t tests indicate no significant difference between the number of photoconverted hair cells at these two timepoints (ant crista p = 0.125, lat crista p = 0.859, utricle p = 0.071). Scale bars = 10 µm. All data is presented as mean ± s.d.

This experiment was repeated for the subsequent week of growth, from 14 to 21 dpf, with again no discernable decrease in photoconverted hair cell number (Fig. S2). Together, these results indicate that little to no hair cell turnover occurs in the zebrafish inner ear organs during larval stages.

### Two hair cell subtypes are added consistently during growth

We wanted to understand how the makeup of sensory organs changes as new hair cells are added. By the larval stage, central and peripheral subtypes exist in the cristae, and striolar and extrastriolar cells are present in the maculae (Qian et al., 2022; Shi et al., 2023; Smith et al., 2020; Smith et al., 2023; Tanimoto et al., 2022). We previously identified marker genes for hair cell subtypes that can be used in Hybridization Chain Reaction Fluorescence in situ Hybridization (HCR-FISH)(Choi et al., 2016; Shi et al., 2023). Here, we used probes against *cabp1b* to label peripheral cells in the cristae. We photoconverted *Tg(myo6b:nls-Eos)* fish at 8 dpf and fixed fish at three subsequent timepoints for imaging: 2 days post-photoconversion (dpp), 7 dpp, and 14 dpp. HCR-FISH was then performed with *cabp1b* probes to distinguish subtypes of “new” (cyan) from “old” (magenta+cyan) hair cells (Fig 4A-C). During this period there is a substantial increase in the number of new hair cells with little change in old hair cells (Fig 4D). Based on the spatial pattern of hair cell addition occurring around the perimeter, we hypothesized that peripheral subtype hair cells would make up the majority of new hair cells. In fact, although *cabp1b+* new hair cells were common at the peripheral poles of the crista, an almost equal percentage of new central-type *cabp1b-* hair cells were added. This even split of new central and peripheral hair cells was consistent at each timepoint examined (Fig 4A-C, E), indicating that both subtypes are added at relatively constant rates. When examining the identity of old hair cells, we observed an increase in the fraction of central to peripheral-type cells over time (Fig. 4A-C, E). This suggests that some hair cells convert from peripheral identity to central identity as sensory patches grow larger in order to maintain the overall ratio of central to peripheral cells.

**Figure 4.**
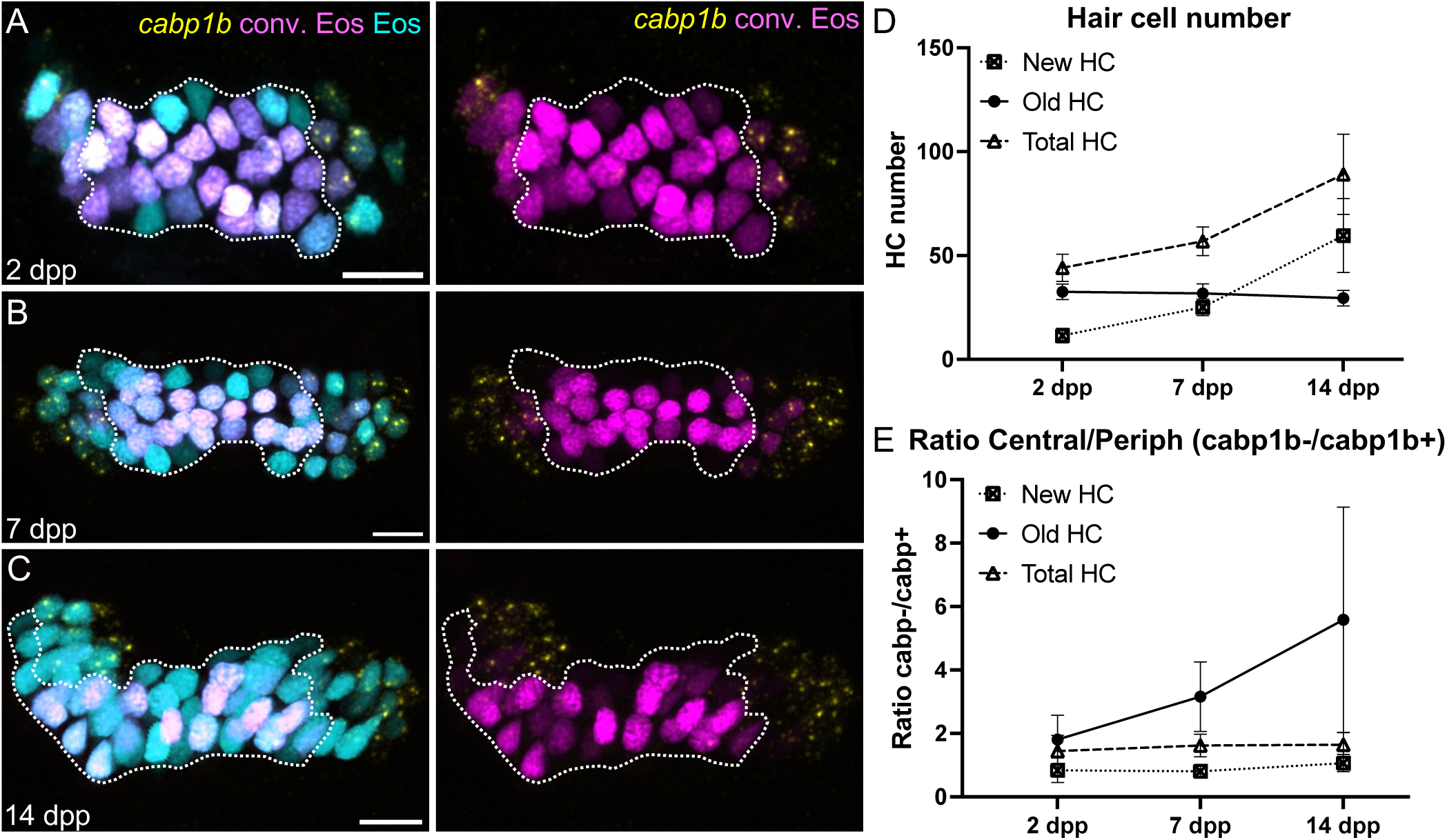
Identification of inner ear hair cell subtypes during larval growth. A-C) Maximum intensity projection images of HCR-FISH probing for *cabp1b* expression in *Tg(myo6b:NLS-Eos)* anterior cristae at A) 2 days post-photoconversion (dpp) (10 dpf, n = 14); B) 7 days dpp (15 dpf, n = 12); and C) 14 dpp (22 dpf, n = 8). Old HC retain photoconverted (magenta) Eos signal while new HC have unconverted (cyan) Eos only. Peripheral-type hair cells are labeled by the *cabp1b* HCR probe (yellow). Dotted outline delineates central, *cabp1b*-region of the sensory patch. Scale bars = 10 µm. D) Increase in hair cell numbers over the course of the experiment. E) ratio of central (*cabp1-*) to peripheral (*cabp1b*+) hair cells over time. The increased ratio for old cells suggests phenotypic conversion from peripheral to central hair cell typeover time. All data is presented as mean ± s.d.

### Crista hair cells are regenerated in the week following ablation

Unlike in the lateral line, hair cells in the inner ear are protected from death caused by immersion in aminoglycoside antibiotics. To overcome this limitation, we designed a *Tg(myo6b:TrpV1-mClover)* transgenic line where the mammalian TRPV1 channel is expressed in target cells (Chen et al., 2016). When exposed to its ligand capsaicin, opening of the mammalian TRPV1 channel results in cell death by cation influx. Endogenous zebrafish Trpv1 is unresponsive to capsaicin, like other non-mammalian forms (Gau et al., 2013). Expressing mammalian TRPV1 under a hair cell specific promoter and exposing the fish to capsaicin results in quick and effective hair cell death in the cristae. We crossed the *Tg(myo6b:TrpV1-mClover)* line to the *Tg(myo6b:nls-Eos)* to identify nascent hair cells. After one hour of treatment, hair cell debris was observed across all three cristae (Fig. 5A-B), and by 4 hours post-treatment this debris had been largely cleared (Fig. 5C). Though this method is highly efficient at killing crista hair cells, hair cell death was inconsistent in the lateral line and was undetectable in the macular organs, likely as a result of different expression levels due to the genetic landscape associated with the location of transgene insertion. Therefore, we focused our subsequent regeneration experiments on the crista. Dose response curves were performed at 5 dpf to determine the appropriate concentration of capsaicin for complete hair cell ablation (Fig. 5D) and found that a 10 μM exposure was sufficient. In all subsequent experiments, larvae were treated with 10 μM capsaicin in system water for one hour. Regeneration experiments were performed in sibling *Tg(myo6b:nls-Eos)* fish in a *nac/roy* background with and without *Tg(myo6b:TrpV1-mClover)*. Due to the relative brightness of Eos, larvae could not be screened for mClover expression under a fluorescent dissecting microscope, even after photoconversion. Thus, fish with dying hair cells were separated from control fish following capsaicin treatment to form the ablated and control groups.

**Figure 5.**
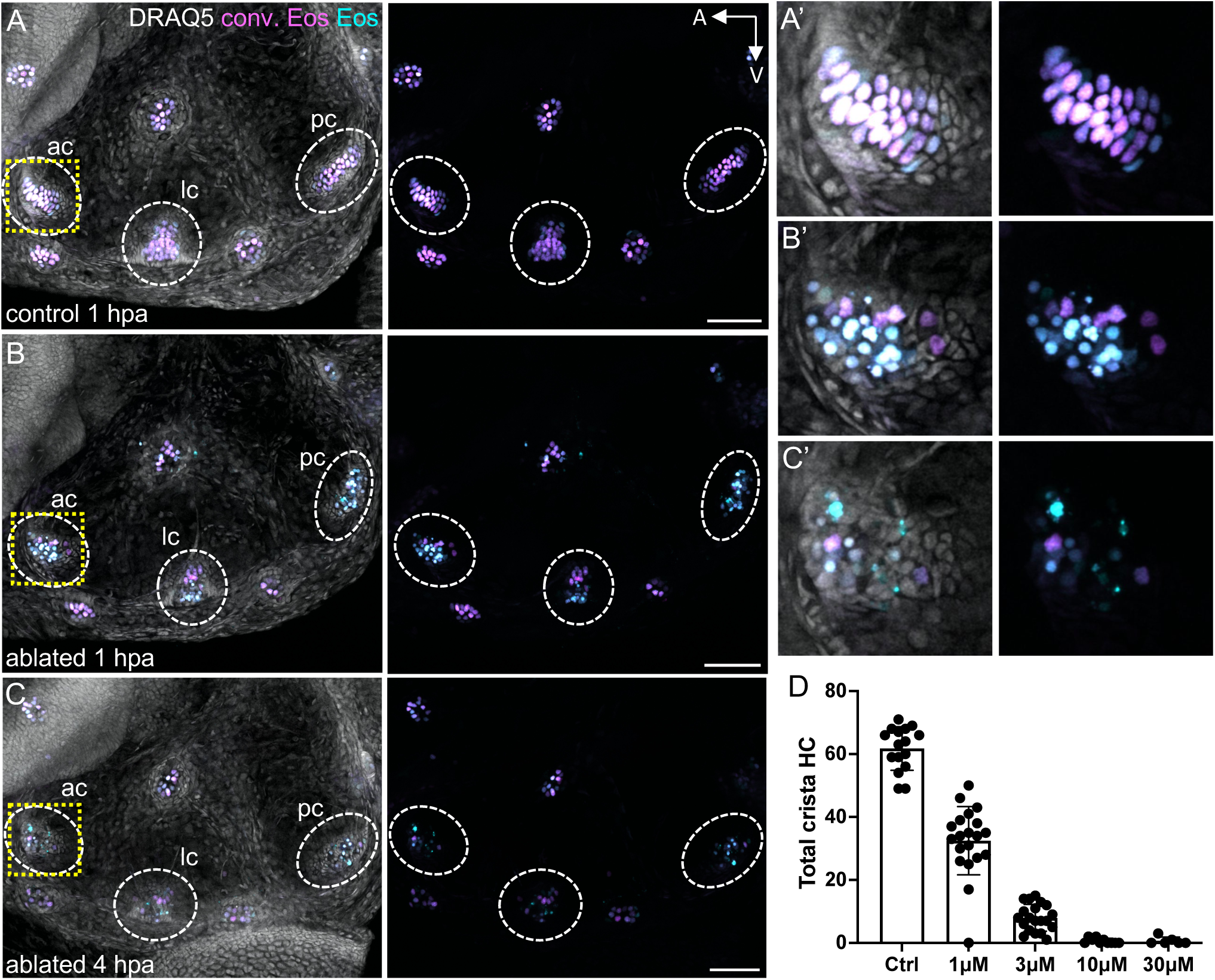
Trpv1-capsaicin hair cell ablation. A) Maximum intensity projection of a photoconverted 8dpf *Tg(myo6b:NLS-Eos)* larval inner ear one hour after capsaicin treatment. B) Maximum intensity projection of a sibling *Tg(myo6b:NLS-Eos);Tg(myo6b:TrpV1-mClover)* inner ear one hour after capsaicin treatment or C) four hours after capsaicin treatment. Images show photoconverted (magenta) and unconverted (cyan) Eos signal with and without DRAQ5-labeled nuclei. Dashed oval regions indicate anterior, lateral, and posterior cristae. Dashed yellow box indicates magnified anterior cristae region shown in A’-C’. D) Dose-response curve for hair cells following one hour of treatment with capsaicin at different concentrations. Control treatment represents DMSO alone. Each data point represents the number of hair cells in combined anterior, lateral, and posterior crista of one fish ear (n = 6-20). All data is presented as mean ± s.d.

To compare hair cell addition following ablation to growth, hair cells were photoconverted and in some fish ablated at 8 dpf, and then fixed at subsequent timepoints to count hair cell nuclei (Fig. 6A). In ablated anterior cristae, the number of new hair cells increased significantly compared to controls over the course of two weeks post-treatment (Fig. 6B-C). Correspondingly, total hair cell number was decreased after capsaicin treatment in ablated fish but slowly recovered to control levels by 14 days post-ablation (dpa) (Fig. 6D). Similar results were obtained for the lateral crista (Fig. S3). No body length difference was observed at any timepoint between control and ablated fish, suggesting that crista hair cell ablation does not affect overall growth rates (Fig. S4). The increased rate of hair cell addition and eventual recovery of hair cell numbers in ablated crista suggest that a regenerative response occurs alongside hair cell addition due to organ growth.

**Figure 6.**
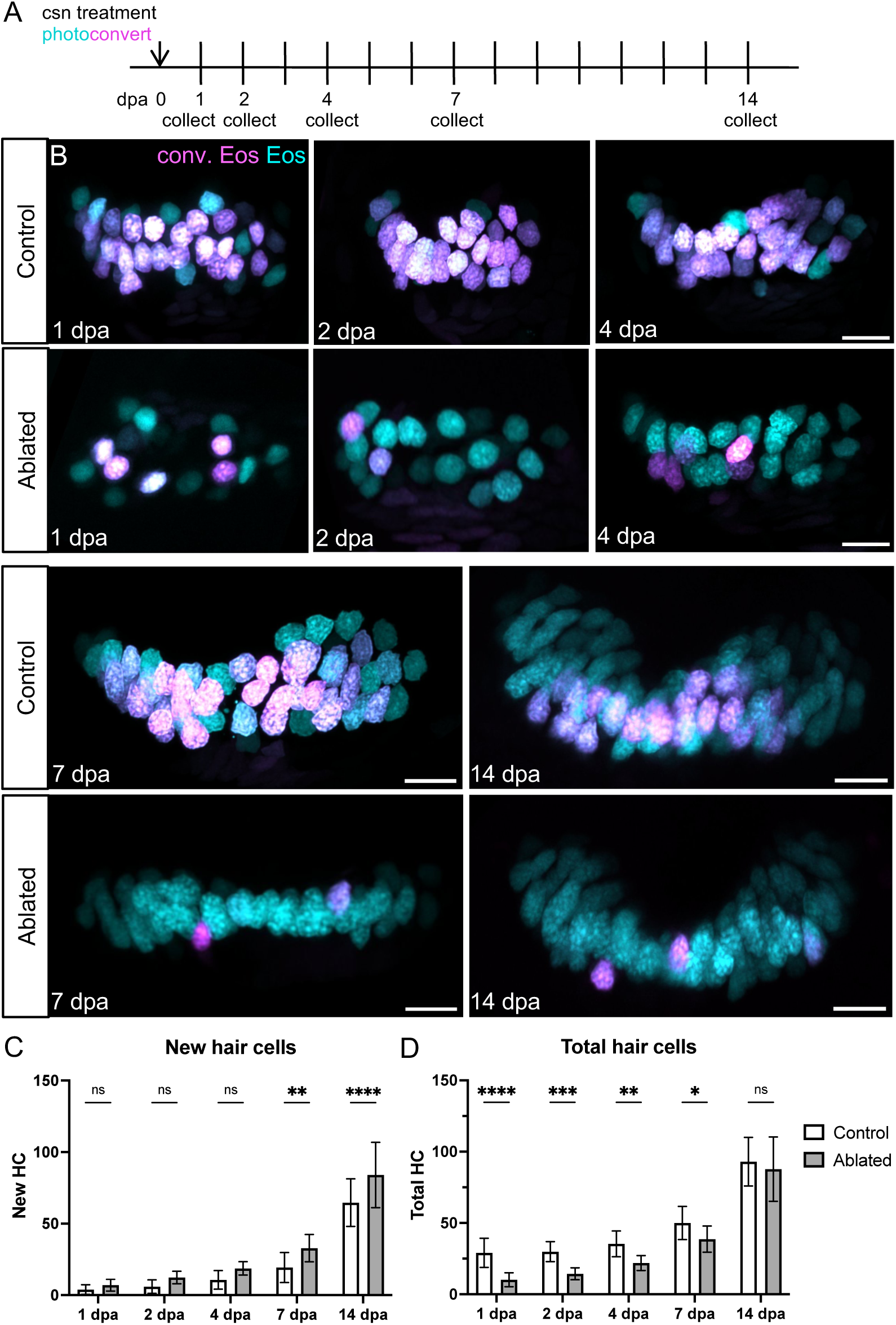
Anterior crista hair cells regenerate during the two weeks following ablation. A) *Tg(myo6b:NLS-Eos)* sibling larvae with or without *Tg(myo6b:TrpV1-mClover)* were photoconverted and treated with capsaicin to ablate hair cells at 8dpf. Larvae were collected at five timepoints over the following two weeks: 1 (n = 22 control, 25 ablated), 2 (n = 13, 20), 4 (n = 19, 18), 7 (n = 16, 13), or 14 (n = 18, 15) days-post ablation. B) Representative maximum intensity projections of anterior crista in control and ablated fish at five timepoints following treatment. Nuclei of cells that survived capsaicin treatment contain photoconverted Eos (magenta). Hair cells newly added after capsaicin treatment have nuclei with only unconverted Eos (cyan). Scale bars = 10 µm. C) Quantification of new (cyan-only) hair cells in ablated and control anterior crista. Two-way ANOVA variation across condition p < 0.0001; Šídák’s multiple comparisons post-hoc test for 7 dpa adjusted p-value = 0.0021, 14 dpa adjusted p-value < 0.0001. D) Quantification of total hair cells in ablated and control anterior crista. Two-way ANOVA variation across condition p < 0.0001; Šídák’s multiple comparisons post-hoc test for 1 dpa adjusted p-value < 0.0001, 2 dpa adjusted p-value = 0.0006, 4 dpa adjusted p-value = 0.0015, 7 dpa adjusted p-value = 0.0342. All data is presented as mean ± s.d.

### Hair cell identity is maintained during regeneration

We next determined whether hair cells regenerated with appropriate spatial identity. We again used HCR probes against *cabp1b* to distinguish peripheral from central type regenerated hair cells. At 2 days post-ablation, newly added *cabp1b*+ and *cabp1b*-hair cells were present in control and ablated conditions (Fig. 7A). In regenerating crista, the percentage of new hair cells of the *cabp1b*+ peripheral type was significantly decreased compared to controls (Fig. 7B). This suggests that the proportion of newly added central-type cells increases in the aftermath of hair cell ablation. To confirm this, we repeated this experiment using HCR probes for *scn5lab*, a marker of central crista hair cells (Fig. S5A). As expected, the proportion of new *scn5lab*+ central-type cells was significantly increased compared to controls (Fig. S5B, C). To determine whether organ patterning returned to that of homeostatic conditions following ablation, we probed for *cabp1b* in 14 dpa fish (Fig. 7C). At this timepoint, when total crista hair cell number in ablated fish has returned to control levels, the overall ratio of central to peripheral hair cells with their regular spatial patterning is also restored (Fig. 7D). Together, these data suggest that a memory of organ patterning and corresponding hair cell identities are maintained in cristae even after extensive hair cell loss.

**Figure 7.**
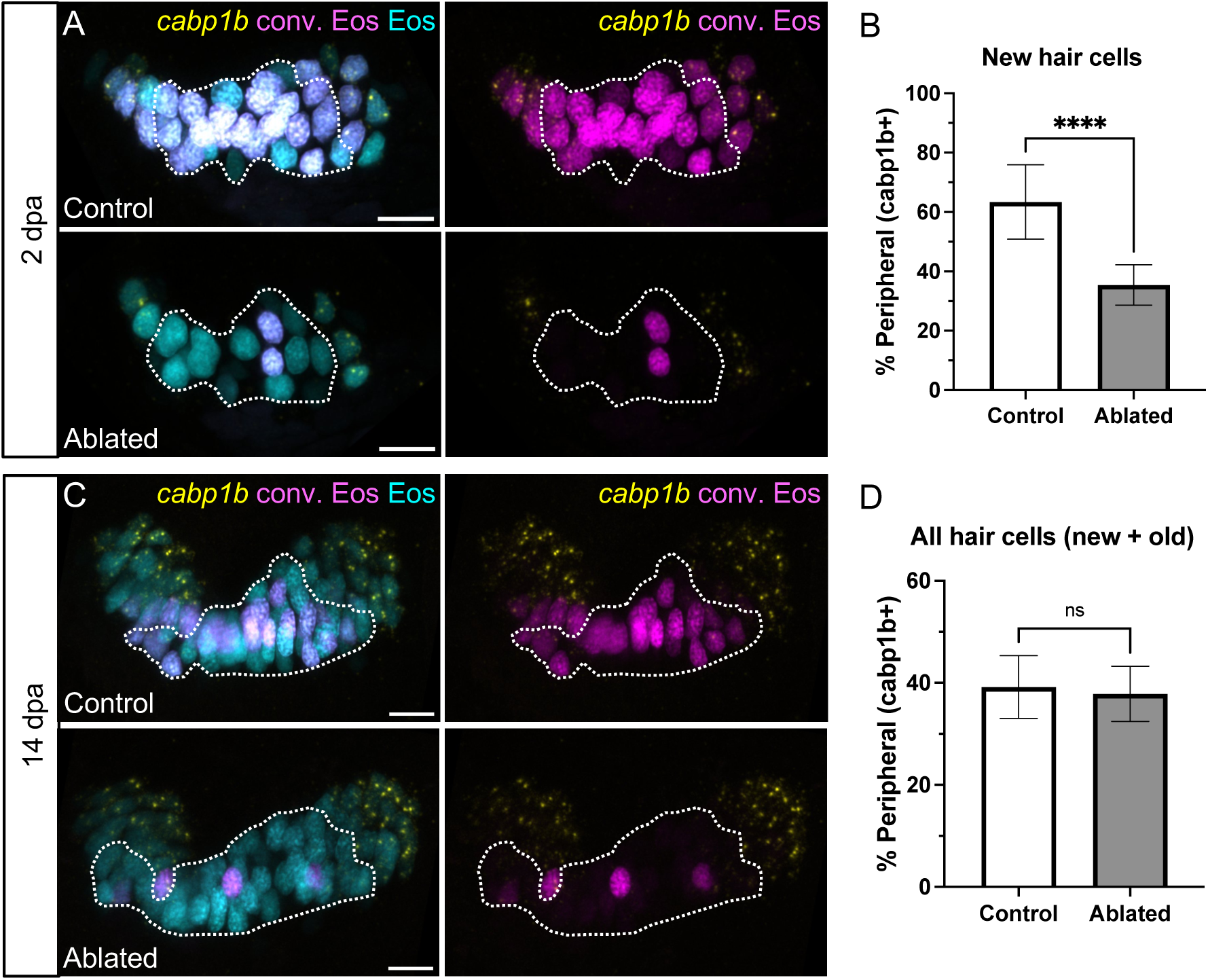
Hair cell central-peripheral patterning is restored following ablation. A) Representative maximum intensity projections of anterior crista in control and ablated fish at 2 dpa with *cabp1b* HCR-FISH. Photoconverted Eos (magenta) and *cabp1b* (yellow) channels are shown with and without unconverted Eos (cyan). Dotted outline delineates central, *cabp1b*-region of the sensory patch. B) Quantification of *cabp1b*+ new hair cells, shown as a percentage of all new (cyan-only) hair cells in control (n = 18) and ablated (n = 16) anterior cristae. Unpaired t test p < 0.0001. C-D) Analogous data to A-B for crista at 14 dpa (n = 18 control, 15 ablated). Unpaired t test p = 0.5226. Scale bars = 10 µm. All data is presented as mean ± s.d.

### Hair cells regenerate primarily by transdifferentiation

To determine whether proliferative mechanisms are used to regenerate hair cells in the zebrafish inner ear, we applied EdU, a thymidine analog that incorporates into the DNA of dividing cells, resulting in labeled daughter nuclei (Salic and Mitchison, 2008). We performed 24-hour EdU pulses in regenerating fish for 0-1 dpa, from 3-4 dpa, and from 6-7 dpa (Fig. 8A). Photoconversion was performed just prior to EdU treatment to identify hair cells added during the EdU pulse. At 1 dpa, EdU labeled hair cells in both control and ablated conditions were rare, less than 1% (Table 1), suggesting that the vast majority of hair cells added immediately post-ablation do not arise from recently dividing progenitors. In both conditions, in cases where rare EdU+ hair cells were observed, they were found paired with an EdU+ supporting cell (Fig. S6), suggesting that a low level of asymmetric division may occur. There was no change at either 4 or 7 dpa, with EdU-labeled hair cells still rare (Table 1), indicating that a later wave of proliferative hair cell regeneration did not occur. We conclude that transdifferentiation is the predominant mechanism by which hair cells are added to regenerating cristae.

**Figure 8.**
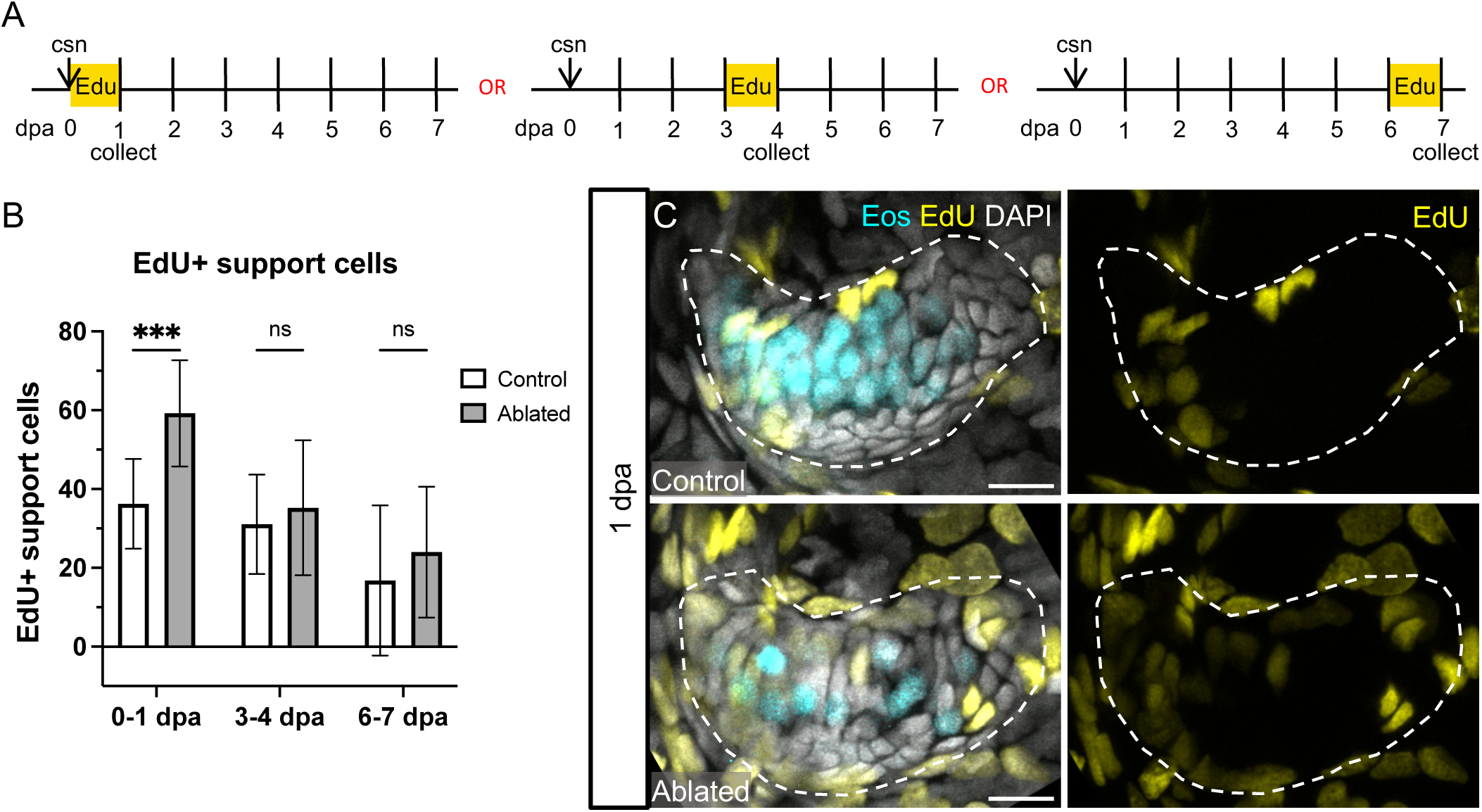
Support cells proliferate in response to hair cell ablation. A) Larvae were incubated in EdU for 24 hours immediately after hair cell ablation, at 3 dpa, or at 6 dpa and collected at the end of the 24-hour incubation. B) Quantification of EdU-labeled support cells in the anterior and lateral cristae combined in control and ablated fish incubated in EdU from 0-1 dpa (n = 13 control, 14 ablated), 3-4 dpa (n = 19, 12), or 6-7 dpa (n = 9, 7). Two-way ANOVA is significant across condition p = 0.0021, Šídák’s multiple comparisons post-hoc test 0-1 dpa adjusted p-value = 0.0004. All data is presented as mean ± s.d. C) Representative maximum intensity projections of anterior crista in control and ablated fish incubated with EdU from 0-1 dpa with Eos-labeled hair cells in cyan and EdU-labeled nuclei in yellow.

**Table 1.** Average EdU+ hair cell counts with percent new hair cells for EdU experiments. Values shown are mean (± s.d.) for combined anterior and lateral cristae.

| Fig. | Timepoint | Control |  |  |  | Ablated |  |  |  |
| --- | --- | --- | --- | --- | --- | --- | --- | --- | --- |
|  |  | EdU+ HC | New HC | Edu % new HC | N | EdU+ HC | New HC | Edu % new HC | N |
| 8 | 0-1 dpa | 0.23 (± 0.48) | - | - | 13 | 0.36 (± 0.63) | - | - | 14 |
| | 3-4 dpa | 0.32 (± 0.48) | 13.26 (± 5.51) | 2.0 (± 3.21) | 19 | 0.42 (± 1.16) | 16.25 (± 5.33) | 1.75 (± 4.86) | 12 |
| | 6-7 dpa | 0.33 (± 0.71) | 14.78 (± 5.52) | 1.80 (± 3.61) | 9 | 0.43 (± 0.53) | 17.29 (± 8.67) | 2.08 (± 2.63) | 7 |
| 9 | 1 dpa | 0.10 (± 0.32) | 14.0 (± 4.71) | 0.40 (± 1.26) | 10 | 0.43 (± 0.53) | 30.57 (± 8.54) | 1.83 (± 2.30) | 7 |
| | 4 dpa | 3.21 (± 1.85) | 29.36 (± 9.25) | 10.8 (± 8.38) | 14 | 5.33 (± 3.74) | 55.0 (± 15.88) | 9.90 (± 6.20) | 9 |
| | 7 dpa | 7.56 (± 4.07) | 74.55 (± 14.53) | 10.01 (± 4.75) | 9 | 12.08 (± 4.35) | 93.23 (± 11.60) | 12.90 (± 4.26) | 13 |

### Hair cell ablation leads to temporary expansion of a supporting cell progenitor pool

In contrast to hair cells, EdU-labeled supporting cells were common at 1 dpa, and significantly more EdU-labeled supporting cells were present compared to controls (Fig. 8B-C). However, there was no significant difference in the number of EdU-labeled supporting cells between control and ablated fish at the 4 or 7 dpa timepoints. These results demonstrate that there is an initial wave of supporting cell proliferation in response to hair cell damage that is not sustained at later periods.

To determine whether supporting cells that divide in response to hair cell ablation ultimately become hair cells, we repeated the regeneration experiment with an EdU pulse during the first 24 hours of regeneration and collected fish at 1, 4 or 7 dpa (Fig. 9A). Again, we observed a significant increase in EdU labeled supporting cells 1 dpa compared to controls (Fig S8), and rare labeled hair cells in both control and ablated conditions (Table 1). EdU-labeled hair cells were increasingly common at the 4 and 7 dpa timepoints in both control and ablated fish (Fig. 9B, C, Table 1). By 7 dpa, significantly more EdU-labeled hair cells were present in ablated crista (Fig. 9B, C), corresponding to the increase in supporting cells labeled at 1 dpa. The total number of new hair cells also significantly increased in ablated compared to control fish (Fig. 9D). When viewed as a percentage of all new hair cells, the fraction of EdU+ hair cells is not significantly different between ablated and control conditions at any timepoint (Fig. 9E). Therefore, supporting cells that divided in response to hair cell ablation are not more likely to differentiate into hair cells. These results suggest that in the wake of hair cell ablation, supporting cells proliferate to increase the progenitor pool, but that this proliferative response is uncoupled to the rate of hair cell differentiation.

**Figure 9.**
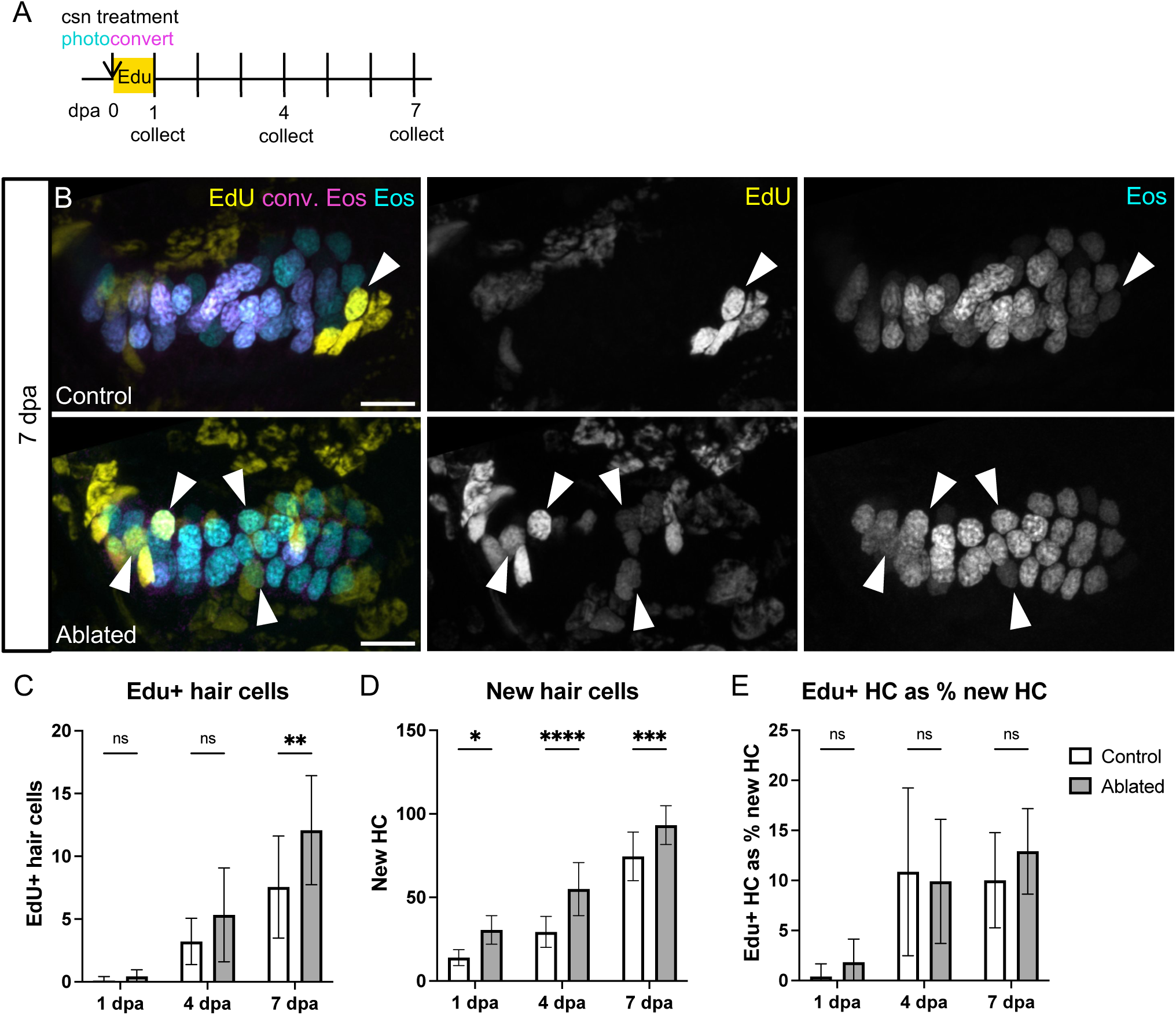
EdU-labeling of hair cells over the week following ablation. A) Larvae were incubated in EdU for 24 hours after photoconversion and hair cell ablation and collected either at the end of the incubation (1 dpa; n = 10 control, 7 ablated) or at 4 (n = 14, 8) or 7 (n = 9, 13) dpa. B) Representative maximum intensity projections of anterior crista in control and ablated fish at 7 dpa. White arrowheads indicate hair cells added since ablation with EdU signal (yellow), unconverted Eos (cyan), and without photoconverted eos (magenta). Scale bars = 10 µm. C) Quantification of EdU+ hair cells in the combined anterior and lateral cristae at each timepoint in control and ablated fish. Two-way ANOVA is significant across condition p = 0.0050, Šídák’s multiple comparisons post-hoc test 7 dpa adjusted p-value = 0.0034. D) Quantification of new (cyan-only) hair cells at each timepoint in control and ablated fish. Two-way ANOVA is significant across condition p < 0.0001, Šídák’s multiple comparisons post-hoc test 1 dpa adjusted p-value = 0.0123, 4 dpa adjusted p-value <0.0001, 7 dpa adjusted p-value = 0.0010. E) EdU+ hair cells as a percentage of new hair cells. Two-way ANOVA with Šídák’s multiple comparisons post-hoc test is not significant across condition at any timepoint. All data is presented as mean ± s.d. combined anterior and lateral cristae.

## DISCUSSION

We describe a steady increase in hair cell number during the growth of inner ear sensory patches during the larval phase of zebrafish development, an approximately month-long period after embryogenesis is complete. We used photoconvertible nuclear-localized Eos to distinguish pre-existing hair cells from newly added hair cells. We found that central and peripheral hair cell subtypes are added at the edges of the organ in a stereotyped pattern based on their location. We document a phenotypic switch of some older hair cells from peripheral to central subtype, presumably to maintain spatial patterning and an overall ratio that slightly favors central-type hair cells. We also found that the number of photoconverted cells in cristae and utricle did not significantly decrease over time, suggesting that there is little hair cell turnover during larval stages.

We provide several lines of evidence that the addition of crista hair cells after damage is more than simply recovery by continued growth. We demonstrate, using photoconversion to parse the timing of differentiation, that new hair cells are added at a faster rate after hair cell kill than during growth. We also found that compared to growth there was an increase in new hair cells of the central subtype, and as a result the organ regenerates the appropriate ratio of subtypes for correct spatial patterning. Finally, there is an increase in supporting cell proliferation in response to hair cell ablation, eventually resulting in more EdU labeled hair cells than under control conditions. If proliferation and hair cell differentiation were directly coupled, we would expect to see a higher proportion of new EdU-labeled hair cells in ablated cristae, as reactive support cells that divided disproportionately regenerated hair cells. The lack of difference between the fraction of EdU labeled new hair cells in control and ablated conditions indicates that the supporting cells dividing in response to ablation are not more likely than others to differentiate into new hair cells. Indeed, our experiments suggest that during growth, supporting cells convert to hair cells using mechanisms uncoupled from cell division, and regenerating hair cells are added through a similar process of transdifferentiation. We hypothesize that the primary regenerative response to damage is to increase the pool of supporting cells available for differentiation into hair cells, employing conversion mechanisms used in growth to add new hair cells. Of particular note, the transient increase in supporting cell proliferation occurs before hair cell replacement, suggesting that the cue for this event is the damage or loss of hair cells rather than depletion of supporting cells through transdifferentiation.

While our current work examines hair cell regeneration in the larval zebrafish cristae over the first month of development, our findings are consistent with previous studies examining regeneration in the zebrafish maculae. Lineage tracing in the embryonic utricle following laser ablation of hair cells provides evidence that supporting cells directly transdifferentiate into nascent hair cells (Millimaki et al., 2010). In the adult saccule, noise damage induces a burst of proliferation that occurs 1-3 days post sound exposure with regenerated hair cell bundles formed in the most damaged area of the organ over approximately 10 days (Schuck and Smith, 2009), a timeline that is consistent with our findings in the cristae. Single-cell RNA-sequencing data from regenerating maculae of adult zebrafish point to the emergence of a transition-state population with qualities of both hair and supporting cells, which could potentially represent actively transdifferentiating cells (Jimenez et al., 2022). Together these studies support a model where damage induces hair cell regeneration through transdifferentiation and expansion of supporting cells through proliferation.

Our findings in the zebrafish inner ear are markedly different from the mechanism of regeneration observed in the zebrafish lateral line system. Following hair cell ablation by ototoxic drug exposure, neuromasts show significant hair cell replacement after 24h and regenerate a full complement of hair cells in just 72 hours (Ma et al., 2008), compared to a gradual replacement of hair cells that we observe in the cristae over the course of two weeks. Lateral line hair cells are regenerated in pairs by symmetrically dividing precursors (Lopez-Schier and Hudspeth, 2006; Mackenzie and Raible, 2012; Romero-Carvajal et al., 2015; Wibowo et al., 2011), while we find those in the cristae are overwhelmingly added by transdifferentiation. The rare examples of EdU-labeled hair cells we observed in the cristae were adjacent to labeled supporting cells, suggesting asymmetric division of precursors.

We speculate these differences may reflect the need in the lateral line system to restore the integrity of organs exposed to the environment on the surface of the fish, while regeneration in the inner ear occurs on top of extensive growth and is needed to restore appropriate spatial patterning rather than organ integrity.

Comparison of growth and regeneration in the inner ear of zebrafish to that in birds reveals both similarities and differences. Regeneration of hair cells in avian auditory and vestibular systems occurs by both transdifferentiation and proliferative replacement. In the regenerating avian utricle, there is evidence that hair cells are replaced both by asymmetric divisions and by transdifferentiation (Scheibinger et al., 2022; Stone et al., 1999). When hair cells are regenerated in the auditory organ, the basilar papilla (BP), they are initially added by wave of transdifferentiation that lasts for several days before a second phase of proliferative hair cell regeneration begins (Roberson et al., 1996; Roberson et al., 2004). To determine whether there is a similar late wave of proliferation in the zebrafish cristae we administered pulses of EdU at timepoints several days after ablation but did not observe any increase in EdU-labeled hair or supporting cells compared to controls. Thus, in the zebrafish larval cristae there appears to be a single mechanism of transdifferentiation for hair cell replacement. In the mature avian vestibule, there is significant hair cell turnover with hair cells having an estimated half-life of about 20-30 days as they are removed and replaced via asymmetric division (Goodyear et al., 1999; Jørgensen and Mathiesen, 1988; Kil et al., 1997; Weisleder and Rubel, 1992). We have observed no evidence for turnover in the zebrafish cristae during larval stages but cannot rule out rare events or turnover at later stages. In the few cases where we observed hair cells labeled by EdU, they were accompanied by with neighboring Edu-labeled supporting cell, suggesting that a small amount of asymmetric division may also occur in the zebrafish inner ear.

Our findings show remarkable similarities to processes that occur in the mammalian vestibular system (Burns et al., 2012a; Wang et al., 2015). When damage is induced in the utricle of neonatal mice, new hair cells are initially generated by transdifferentiation of supporting cells, with an accompanying wave of supporting cell proliferation detected by EdU incorporation. In the following weeks a fraction of these EdU-labeled cells become new hair cells. However, the regenerative response is greatly diminished after the first week postpartum. These regenerative events parallel processes that occur during the normal postnatal growth of the mouse utricle, where approximately half of hair cells are added over the three weeks after birth from supporting cells that last divided before birth (Burns et al., 2012b). In adult mice, limited regeneration occurs by transdifferentiation of supporting cells with no detected proliferative response for their replacement, and as a consequence an overall reduction in supporting cell numbers is observed (Golub et al., 2012). Hair cell turnover, while detectable in the adult mouse utricle, is rare and not associated with supporting cell proliferation (Bucks et al., 2017). Taken together these studies support the idea that there is uncoupled potential for both proliferative and transdifferentiation responses in the mouse utricle that wane over time.

Our study establishes the zebrafish inner ear as a model for hair cell regeneration that parallels processes that are functional for a limited period in mammals. A major difference between mammals and zebrafish is that they lose their ability to regenerate in response to damage (Burns et al., 2012a; Cox et al., 2014) even in response to exogenous factors such as altering Notch signaling or inducing Atoh1 expression (Liu et al., 2012; Maass et al., 2015). Whether mammals lose their ability to regenerate due to epigenetic changes affecting chromatin accessibility (Tao et al., 2021), alterations in cell cycle regulation (White et al., 2006), changes in tissue architecture (Burns and Corwin, 2014; Collado et al., 2011) or a combination with other unknown factors remains an area of active study.

Zebrafish have a remarkable ability to regenerate many organs, including the heart, liver, kidney, fin, retina, and central nervous system (reviewed in Marques et al., 2019), some of which show similarities to the inner ear regeneration mechanism we describe here. In the zebrafish olfactory bulb, death of sensory neurons by chemical exposure results in proliferation of the precursor pool during the first 24 hours following neuron death (Ma et al., 2018). Transdifferentiation has been observed during regeneration of other zebrafish organ systems, particularly in response to severe organ damage. After major damage to the liver, biliary epithelial cells proliferate and transdifferentiate into regenerated hepatocytes (Choi et al., 2014). In the zebrafish pancreas, upon ablation of insulin-responsive β-cells, some Ill-cells transdifferentiate into β-cells while others respond by proliferating, presumably to replace converting Ill-cells (Ye et al., 2015). Other organs do not exhibit transdifferentiation but rely on a resident population of multipotent cells that act in growth and regeneration. Like the in the ear and other organs, zebrafish kidneys grow throughout life in proportion to fish size (Zhou et al., 2010). Some ototoxic drugs, such as aminoglycoside antibiotics, also demonstrate nephrotoxicity. After injection of the aminoglycoside gentamicin, adult zebrafish regenerate nephrons over the course of two weeks (Diep et al., 2011). In this case, regeneration is facilitated by a resident stem cell population that acts both in adult nephrogenesis as well as regeneration (Diep et al., 2011). In the adult zebrafish central nervous system, the telencephalon contains radial glia that proliferate under homeostatic conditions (Rothenaigner et al., 2011). These same glia respond to lesion injury with proliferation and give rise to neuroblasts that migrate to the site of injury where they differentiate into mature neurons (Kroehne et al., 2011). Our work indicates that support cells of the inner ear may represent a similar resident facultative progenitor population that can self-renew and generate hair cells during growth and regeneration. Whether inner ear support cells are comprised of subpopulations with differential potential to give rise to hair cells remains an unanswered question.

## METHODS

### Fish maintenance

Experiments were conducted on larval zebrafish between 5dpf and approximately 45dpf (up to 11.0mm SL). Larvae were raised in E3 embryo medium (14.97 mM NaCl, 500 mM KCL, 42 mM Na2HPO4, 150 mM KH2PO4, 1 mM CaCl2 dihydrate, 1 mM MgSO4, 0.714 mM NaHCO3, pH 7.2) at 28.5°C and placed on the nursery system at 5dpf. All transgenic fish lines were crossed into a *nac/roy* background (Lister et al., 1999; Ren et al., 2002; White et al., 2008) to facilitate inner ear imaging. During experiments, larval fish were returned to the nursery system between treatment and collection timepoints, except during EdU incubation or when collected immediately after treatment. Zebrafish experiments and husbandry followed standard protocols in accordance with University of Washington Institutional Animal Care and Use Committee guidelines.

### Photoconversion

Larvae were transferred to a 60 x 15mm petri dish and placed in a freezer box lined with aluminum foil. An iLumen 8 UV flashlight (procured from Amazon) was affixed in the freezer box lid and positioned over the dish. Larvae were exposed to UV light for 10 min before being returned to standard 100 x 15mm petri dishes to await experimentation.

### TrpV1 hair cell ablation

Capsaicin (Sigma-Aldrich, #M2028) was resuspended in DMSO and stored at -20°C until use. Dose-response curves were performed on *Tg(myo6b:TrpV1-mClover)* in both *AB and Nac/Roy backgrounds. There were no apparent differences in response to capsaicin treatment between fish of the two backgrounds. 10 μM capsaicin was determined to be an appropriate dose to effectively ablate cristae hair cells when treated for 1 hour at 28.5°C. The brightness of Eos in the *Tg(myo6b:nls-Eos)* line prevents normal fluorescent dissecting scope screening for *Tg(myo6b:TrpV1-mClover)*, even after Eos has been photoconverted. 8dpf *Tg(myo6b:nls-Eos)* siblings with and without *Tg(myo6b:TrpV1-mClover)* were treated with 10 μM capsaicin for one hour at 28.5°C. Larvae were washed 3 x 5 minutes in system water. Larvae were then screened for dying hair cells to indicate the presence (ablated) or absence (control) of *TrpV1-mClover*. Ablated and control fish were separately returned to the nursery system to await collection.

### EdU treatment and visualization

Larvae were incubated in 500μM F-ara-EdU (Sigma, #T511293) for 24 hours at 28.5°C. Click-iT protocol was modified from Salic and Mitchison, 2008. Briefly, larvae were fixed in 4% paraformaldehyde at 4°C for 18-48 hours, depending on their size, then washed with PBS containing 0.1% Tween20 for 3 x 10 minutes. Larvae were permeabilized in 0.5% TritonX-100 in PBS for 30 minutes and washed 3 x 10 minutes with PBS alone. Reaction solution was prepared fresh each time: 2 mM CuSO_4_, 10 mM Alexa Fluor 647 azide, and 20 mM sodium ascorbate in PBS. Fish were incubated in reaction solution for 1 hour in the dark at room temperature, washed 3 x 20 minutes with PBS, and stored in the dark at 4°C until imaging.

### HCR FISH

Hybridization chain reaction in situ hybridizations (Molecular Instruments, HCR v3.0) were performed as directed for whole-mount zebrafish embryos and larvae (Choi et al., 2016; Choi et al., 2018). Briefly, larvae were fixed in 4% PFA at 4°C for 18-48 hours. Larvae were washed with PBS and transferred to MeOH to be stored at -20°C until use. Larvae were rehydrated using a gradation of MeOH and PBST washes, treated with proteinase K for 25 minutes and post-fixed with 4% PFA for 20 minutes at room temperature. For the detection phase, larvae were pre-hybridized with a probe hybridization buffer for 30 minutes at 37 °C, then incubated with probes overnight at 37°C. Larvae were washed with 5X SSCT to remove excess probes. For the amplification stage, larvae were pre-incubated with an amplification buffer for 30 minutes at room temperature and incubated with hairpins overnight in the dark at room temperature. Excess hair pins were removed by washing with 5X SSCT. Larvae were transferred to storage buffer and kept in the dark at 4°C until imaging.

### Fixation and imaging preparation

Larvae were fixed in 4% paraformaldehyde at 4°C for 18-48 hours, depending on their size. Larvae were washed 3 x 15 mins in PBS containing 0.1% Tween20 and transferred to storage buffer (PBS containing 0.2% Triton, 1% DMSO, 0.02% sodium azide, and 0.2% BSA). Samples were stored for no more than 3 weeks at 4°C before imaging. Fixed fish were mounted by first drawing a thin ring of vacuum grease on the underside of a coverslip. One or more specimens were placed on their side in the center of the ring along with 1-2 drops of PBS or other storage solution. A second coverslip was placed on top and gently pushed down at the sides to create a seal around the samples to prevent evaporation and drifting while imaging. Coverslip “sandwiches” were overlayed on a flat ruler under a dissecting microscope, and standard length for each fish was measured, estimating to the nearest 0.25mm. Hair cells in the cristae and utricle were counted in fixed, intact fish as much as possible. When larval fish grew beyond approximately 8 mm, it became necessary to dissect the ear in order to image and perform accurate hair cell counts.

### Imaging

Images for development, turnover, and regeneration timeline experiments were captured on a Zeiss LSM-880 with Airyscan 1.0 functionality. Z-stacks of inner ear organs were taken using a 20X air objective at intervals of 0.32 μm. Development, turnover, regeneration timeline, EdU, and HCR experimental images were captured using a Zeiss LSM-980 with Airyscan 2.0. Z-stacks of inner ear organs were taken using a 20X water objective at intervals of 0.58 μm. Z-stacks of whole ears were taken using a 10X air objective at intervals of 1.32 μm. All Airyscan processing was performed at standard strength using Zen Blue software (Zeiss, www.zeiss.com). Image processing and data analysis were performed using Fiji (Schindelin et al., 2012).

### Statistical analysis

Power analyses were performed in G*Power (Faul et al., 2007) using preliminary data to determine sample sizes. All other statistical analyses were performed in GraphPad Prism version 10.1.0 (GraphPad Software, Boston, Massachusetts USA, www.graphpad.com).

### DATA AVAILABILITY

All data available upon request to corresponding author.

## ACKNOWLEDGEMENTS

We would like to thank David White, Jessica Knight, George Sanders DVM, and the rest of the University of Washington zebrafish facility for fish care. We thank Tor Linbo and Brenna N. Linton for assistance with breeding and screening of fish.

## AUTHOR CONTRIBUTIONS

Marielle O. Beaulieu, Conceptualization, Methodology, Validation, Investigation, Writing - original draft, Writing – review and editing, Visualization, Funding acquisition; Eric D. Thomas, Methodology; David W. Raible, Conceptualization, Methodology, Writing – review and editing, Supervision, Funding acquisition

---

**Figure S1:**
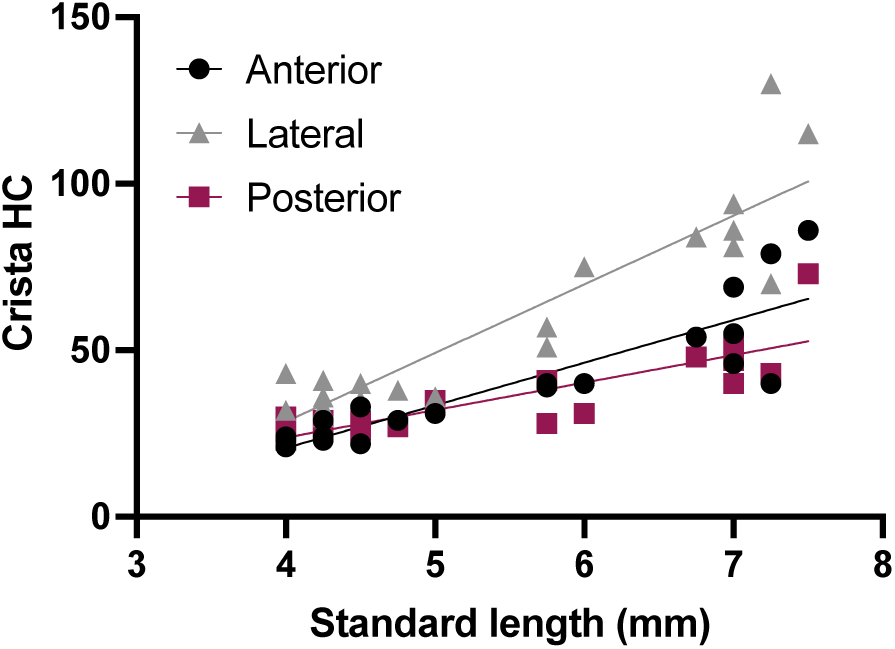
The posterior crista is similar in size to the anterior crista. Hair cell counts from the anterior, lateral, and posterior cristae from the same set of fish. Each data point represents one ear from one fish (n = 21). Linear regression of anterior crista slope 12.8 = ± 1.59, R^2^ = 0.773; lateral crista slope = 20.61 ± 2.12, R^2^ = 0.832; posterior crista slope = 8.26 ± 1.21, R^2^ = 0.720.

**Figure S2:**
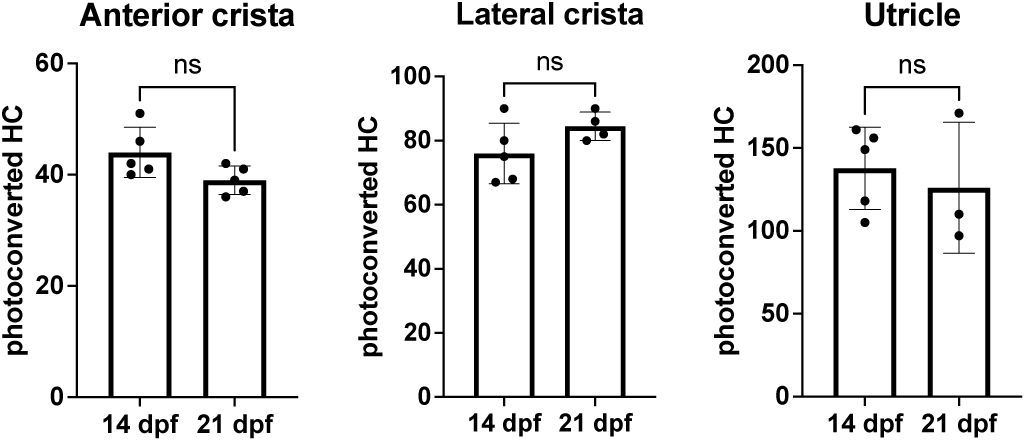
Little hair cell turnover occurs in the third week post-fertilization. Quantification of anterior crista, lateral crista, and utricle photoconverted hair cells at 14 (n = 5 ant crista, 5 lat crista, 5 utricle) and 21 dpf (n = 5 ant crista, 4 lat crista, 3 utricle). Mann-Whitney tests indicate no significant difference between the number of photoconverted hair cells at these two timepoints (ant crista p = 0.095, lat crista p = 0.174, utricle p = 0.786). All data is presented as mean ± s.d.

**Figure S3:**
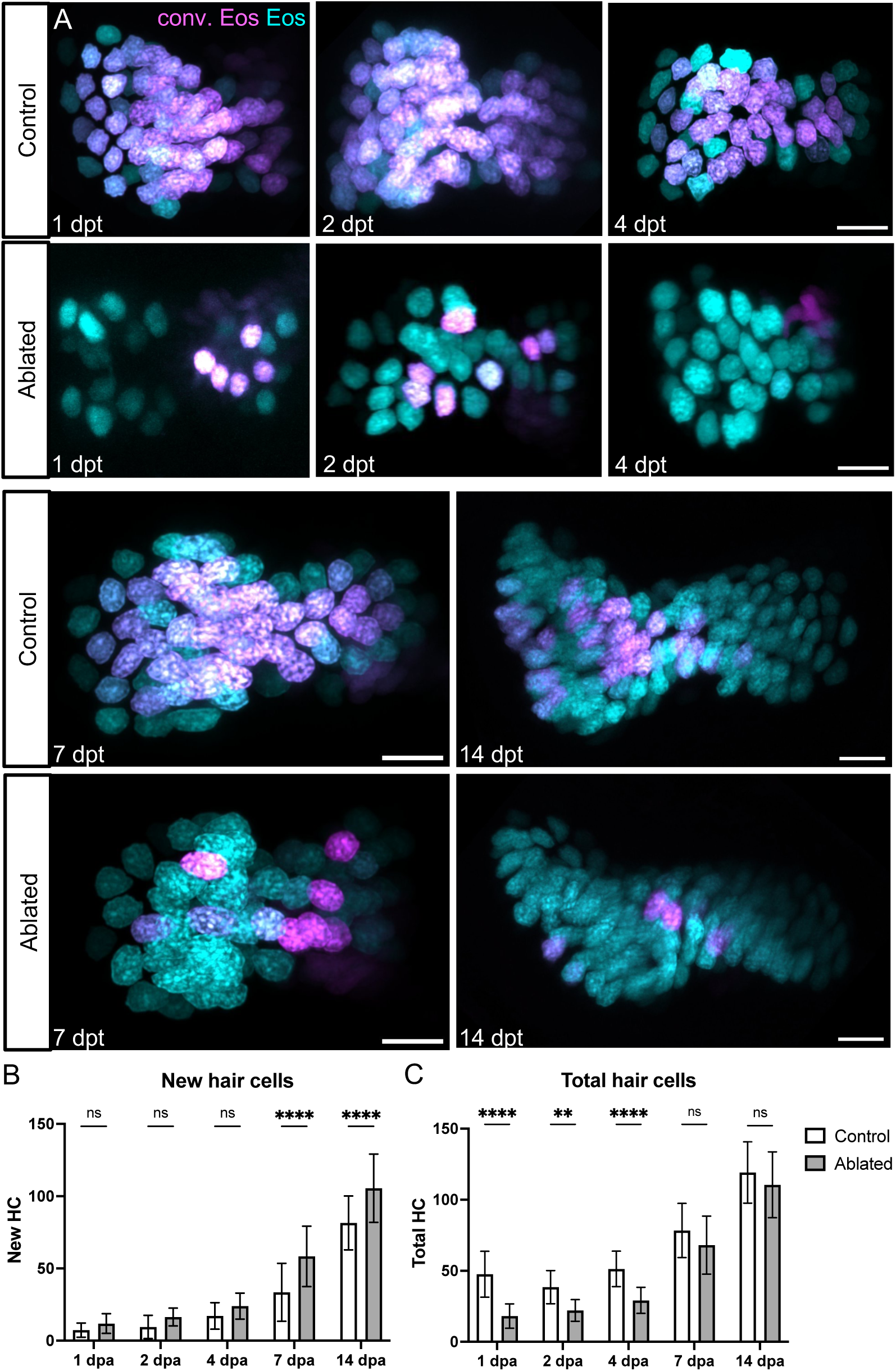
Lateral crista hair cells regenerate during the two weeks following ablation. A) *Tg(myo6b:NLS-Eos)* sibling larvae with or without *Tg(myo6b:TrpV1-mClover)* were photoconverted and treated with capsaicin to ablate hair cells at 8dpf. Larvae were collected at five timepoints over the following two weeks: 1 (n = 22 control, 25 ablated), 2 (n = 20, 22), 4 (n = 19, 17), 7 (n = 16, 12), or 14 (n = 18, 14) days post-ablation (dpa). Representative maximum intensity projections of lateral crista in control and ablated fish at five timepoints following treatment. Nuclei of cells that survived capsaicin treatment contain photoconverted Eos (magenta). Hair cells newly added after capsaicin treatment have nuclei with only unconverted Eos (cyan). Scale bars = 10 µm. B) Quantification of new (cyan-only) hair cells in ablated and control lateral crista. Two-way ANOVA variation across condition p < 0.0001; Šídák’s multiple comparisons post-hoc test for 7 dpa adjusted p-value < 0.0001, 14 dpa adjusted p-value < 0.0001. C) Quantification of total hair cells in ablated and control anterior crista. Two-way ANOVA variation across condition p < 0.0001; Šídák’s multiple comparisons post-hoc test for 1 dpa adjusted p-value < 0.0001, 2 dpa adjusted p-value = 0.0051, 4 dpa adjusted p-value < 0.0001. All data is presented as mean ± s.d.

**Figure S4:**
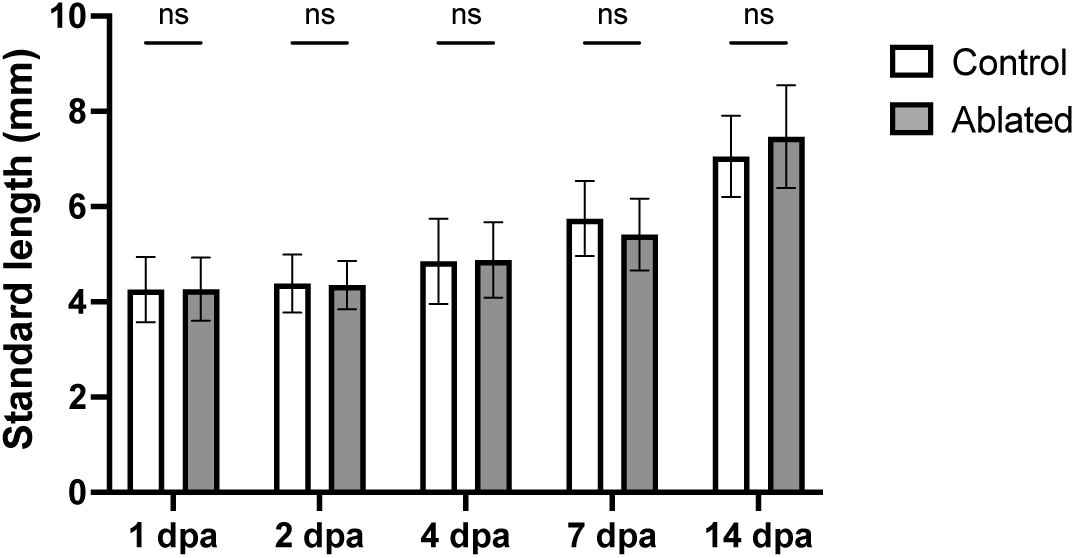
Crista hair cell ablation does not affect larval growth. Standard length measurements across 1 (n = 23 control, 27 ablated), 2 (n = 13, 22), 4 (n = 12, 17), 7 (n = 15, 6), or 14 (n = 32, 16) dpa timepoints for ablated and control larvae. Two-way ANOVA with Šídák’s multiple comparisons post-hoc test indicates no significant differences across condition at any timepoint.

**Figure S5:**
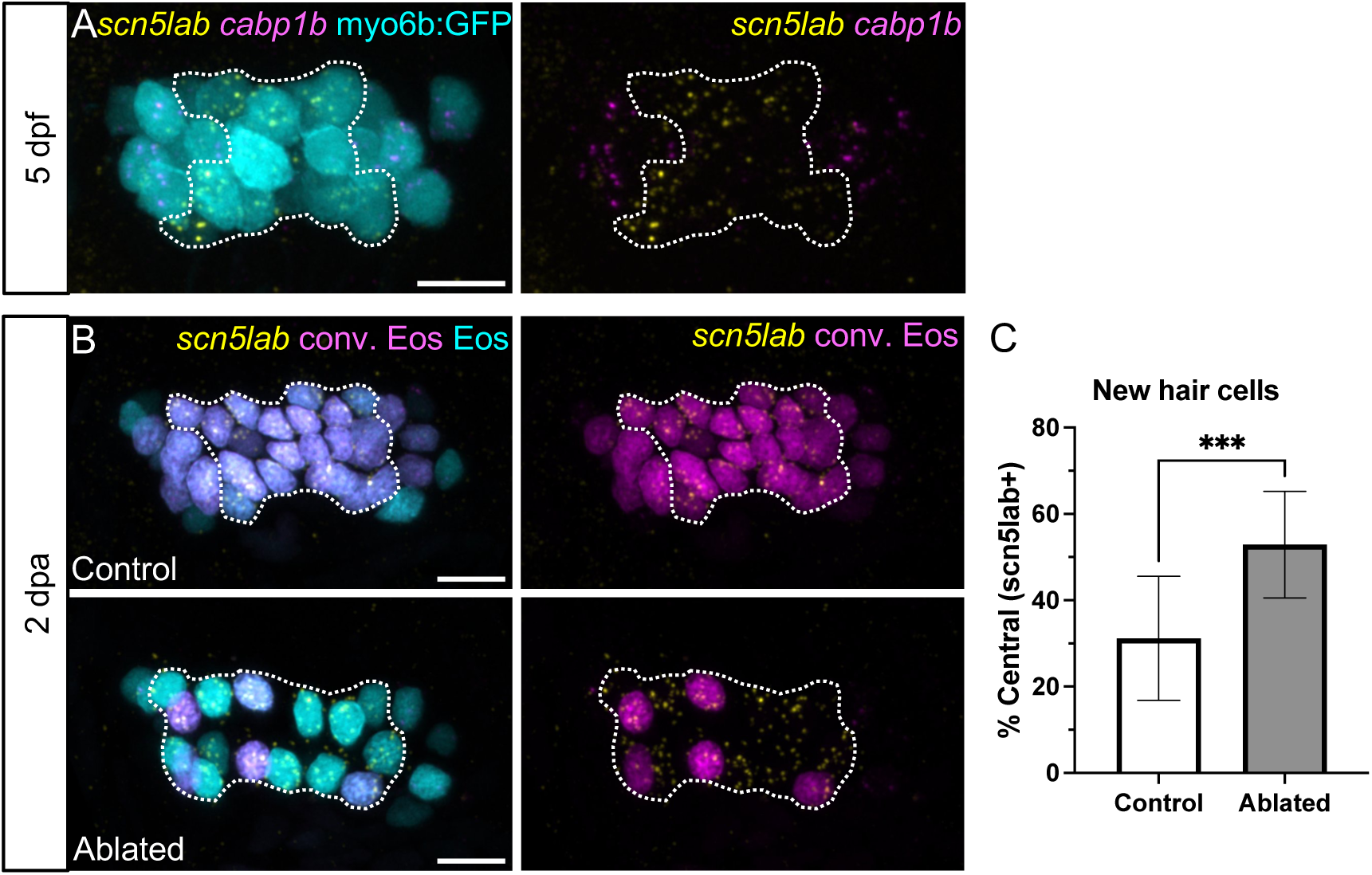
Central-type hair cells are preferentially added following hair cell ablation. A) Representative maximum intensity projections of the anterior crista of a *Tg(myo6b:GFP)* (cyan) 5 dpf larvae treated with HCR *cabp1b* (magenta) and *scn5lab* (yellow) probes to label peripheral- and central-type hair cells, respectively. Dotted outline delineates central, *cabp1b*-;*scn5lab*+ region of the sensory patch. B) Representative maximum intensity projections of anterior crista in control and ablated fish at 2 dpa with *scn5lab* HCR-FISH. Photoconverted Eos (magenta) and *scn5lab* (yellow) channels are shown with and without unconverted Eos (cyan). Dotted outline delineates central, *scn5lab*+ region of the sensory patch. Scale bars = 10 µm C) Quantification of *scn5lab*+ new hair cells, shown as a percentage of all new (cyan-only) hair cells in control (n = 13) and ablated (n = 17) anterior cristae. Unpaired t test p = 0.0001.

**Figure S6:**
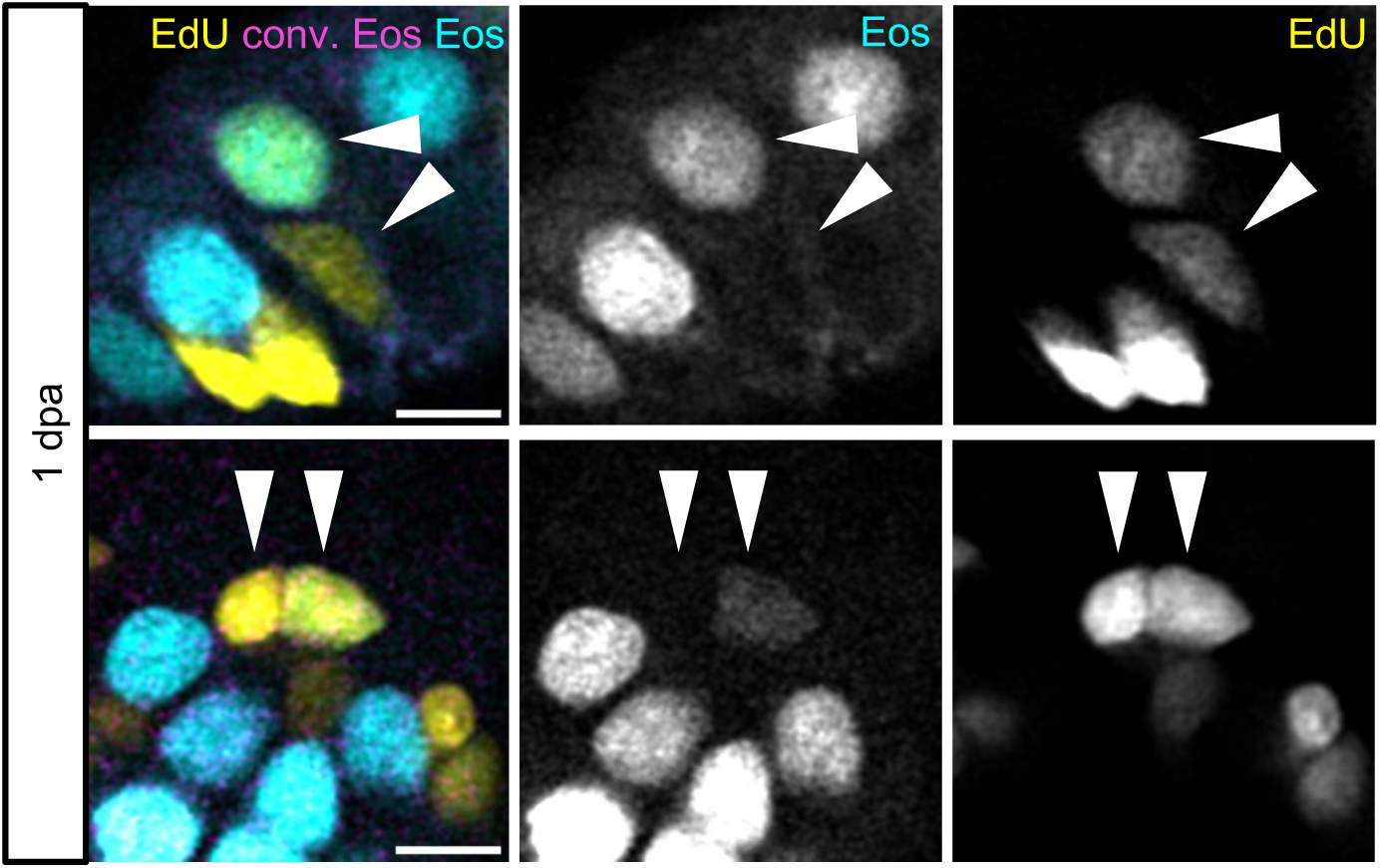
EdU-labeled hair cell-supporting cell pairs are observed following hair cell ablation. Two examples of hair cell-supporting cell EdU+ pairs in ablated fish after 24h EdU incubation (1 dpa). Arrows indicate pairs where both cells are labeled with EdU (yellow), but only one expresses the Eos (cyan) hair cell marker. Scale bars = 5 µm

**Figure S7:**
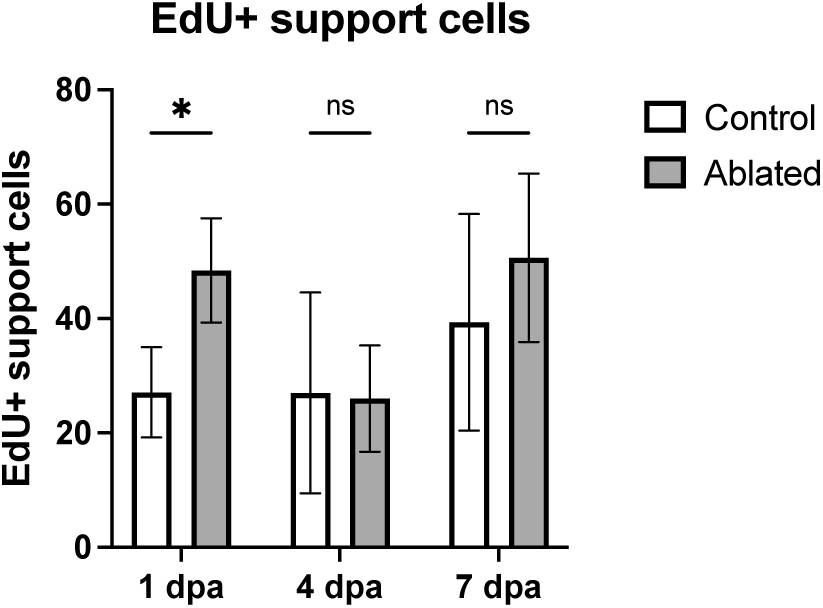
EdU-labeling of supporting cells over the week following ablation. Larvae were incubated in EdU for 24 hours after photoconversion and hair cell ablation and collected either at the end of the incubation (1 dpa; n = 10 control, 7 ablated) or at 4 (n = 14, 8) or 7 (n = 9, 13) dpa. Quantification of EdU+ supporting cells in the combined anterior and lateral cristae at each timepoint in control and ablated fish. Two-way ANOVA is significant across condition p = 0.0069, Šídák’s multiple comparisons post-hoc test 1 dpa adjusted p-value = 0.0107. All data is presented as mean ± s.d.

